# Genotype-Dependent Dysregulation of the *MDM2–p53* Axis and Breast Cancer Susceptibility in Bangladeshi Women: A Cas-Control Study

**DOI:** 10.64898/2026.05.18.726100

**Authors:** Mahim Hassan Chowdhury, Farzana Islam, Adiba Ayesha Khan, Md. Ayman Siddique, Nushaiba Binte Hasan, Mahinul Islam Samrat, Mehrin Haque Tanisha, Jarin Tasnim, Samiha Mahjabin, Md. Nazmul Islam, Md. Aminul Haque

**Author notes:** Equal contribution (Mahim Hassan Chowdhury, Farzana Islam). **Emails** Mahim Hassan Chowdhury, Farzana Islam, Adiba Ayesha Khan, Md. Ayman Siddique, Nushaiba Binte Hasan, Mahinul Islam Samrat, Mehrin Haque Tanisha, Jarin Tasnim, Samiha Mahjabin, Md. Nazmul Islam. Corresponding Author **Md. Aminul Haque**.

## Abstract

**Background:** The *MDM2–p53* signaling pathway plays a central role in tumor suppression, and genetic variants that disrupt this pathway may influence breast cancer (BC) susceptibility. However, data from South Asian populations, particularly Bangladesh, remain limited.

**Methods:** A case–control study was conducted in Bangladeshi women, including BC patients and healthy controls (HCs). Genotyping of *MDM2* polymorphisms was performed using PCR-based methods. Circulating *MDM2* and *p53* protein levels were measured using enzyme-linked immunosorbent assays (ELISA). Associations between genotype, protein levels, BC status, and clinicopathological features were evaluated using appropriate statistical models.

**Results:** A strong and genotype-specific association was observed for *MDM2* rs2279744. Women carrying the heterozygous TG genotype had a markedly increased risk of BC across additive, dominant, and over-dominant models, whereas the GG genotype showed a protective effect under the recessive model. In contrast, rs937282 did not show a significant association with BC risk. Circulating *MDM2* levels were significantly elevated in patients compared with controls and varied by rs2279744 genotype, while circulating *p53* levels showed an opposite trend. A strong inverse correlation was observed between serum *MDM2* and *p53* levels, supporting dysregulation of the *MDM2–p53* feedback loop. Elevated *MDM2* levels were also noted in *HER2-*positive and triple-positive BC subtypes.

**Conclusion:** Together, these findings indicate that the *MDM2* rs2279744 polymorphism contributes to BC susceptibility in a genotype-specific manner, likely through disruption of the *MDM2–p53* regulatory balance. However, the absence of functional validation limits direct causal inference.

## 1. Introduction

BC is one of the most common types of cancer and the main cause of cancer-related death in women around the world [1]. Every year, more than two million new cases are diagnosed [2, 3]. Breast and cervical cancers make up almost 40% of all female cancers in Bangladesh [4, 5]. This is mostly because they are diagnosed late and there are few screening options [6]. This shows how important it is to find genetic risk factors. Genetic factors are very important when it comes to breast cancer risk, even though lifestyle and reproductive factors also play a role [7, 8]. In addition to high-risk mutations like *BRCA1* and *BRCA2*, prevalent single nucleotide polymorphisms may modify gene regulation in critical pathways governing cell cycle progression, DNA repair, and apoptosis, thereby increasing susceptibility to breast cancer [8, 9].

Mouse double minute 2 *(MDM2*) is a major controller of the tumor suppressor *p53*. It keeps the genome stable by controlling cell-cycle arrest, DNA repair, and apoptosis [10–12]. *MDM2* normally keeps *p53* from working too hard, but when it is overexpressed, it can stop *p53* from working and help genetically unstable cells survive, which can lead to cancer [13–18].

Regulatory single nucleotide polymorphisms (SNPs), especially those in promoter and untranslated regions, usually change how much a gene is expressed instead of turning it on or off [19]. These kinds of variants can change transcriptional dynamics, allelic balance, and regulatory noise, which can change signaling pathways in a non-linear way that depends on the genotype [20]. The *p53–MDM2* feedback loop is very tightly controlled, so even small changes in *MDM2* expression can have a big effect on *p53* activity. This means that heterozygous and homozygous variants may have different biological effects [15].

There are two SNPs in *MDM2* that are very interesting. The first one, rs2279744 (also known as SNP309, T>G), is in the promoter region of the gene [21]. The G allele enhances the binding affinity for the transcription factor Sp1, leading to elevated *MDM2* expression and diminished *p53* activity [22, 23]. Numerous studies conducted in Asian and European populations have demonstrated that this variant correlates with an increased risk of various cancers, including breast cancer. The magnitude and orientation of this risk, however, fluctuate based on ethnicity and study design [24, 25]. The second variant, rs937282 (C>G), is found in the 5′ untranslated region (5′UTR) of *MDM2* [26]. This area is important for controlling the stability and efficiency of mRNA translation. Consequently, modifications in this context may affect *MDM2* protein concentrations and impact tumor biology [27]. Some studies have found that rs937282 may raise the risk of cancer, while others have found little or no link. This shows that more research is needed for specific populations [28–30].

In addition to genetic susceptibility, the levels of *MDM2* and *p53* proteins in the blood have been found to be possible markers of *p53* pathway dysregulation [31]. In numerous malignancies, including breast cancer, elevated *MDM2* levels and altered *p53* concentrations in serum have been documented [32]. Nonetheless, the correlation among *MDM2* polymorphisms, circulating protein concentrations, and tumor biological traits, including hormone receptor status, is not fully elucidated. Breast cancer is a diverse disease characterized by distinct molecular subtypes delineated by estrogen receptor (*ER*), progesterone receptor (*PR*), and human epidermal growth factor receptor 2 (*HER2*) status [33]. New evidence suggests that the way oncogenic signaling pathways and the *p53–MDM2* axis interact may be different in receptor-defined subtypes [34].

Due to the scarcity of data from South Asian populations, especially Bangladesh, this study sought to examine the correlation between two regulatory *MDM2* polymorphisms (rs2279744 and rs937282) and breast cancer susceptibility in Bangladeshi women. We also looked at how circulating levels of *MDM2* and *p53* vary depending on genotype, how they relate to different types of BC receptors, and how *MDM2* and *p53* relate to each other as a sign of pathway dysregulation. This study aims to enhance the understanding of how the disruption of the *MDM2– p53* association may influence breast cancer risk in this population by integrating genetic, molecular, and clinical data.

## 2. Materials and Methods

### 2.1. Study Subjects

All participants underwent thorough physical, neurological, and laboratory evaluations to confirm the absence of any significant physical or neurological conditions. HCs and BC patients were recruited from Dhaka, Bangladesh. The study included 112 women with BC and 124 age-sex matched HCs, enrolled between April 1, 2023, and June 30, 2023. This was an exploratory pilot study; therefore, the sample size was determined by feasibility and availability of eligible subjects during the study period, and an a priori power calculation was not performed. BC patients were recruited from National Cancer Research Institute Hospital in Mohakhali, Dhaka, utilizing well-equipped inpatient and outpatient facilities for assessments. A qualified oncologist diagnosed BC patients using standard diagnostic criteria and evaluated the HCs. Demographic and clinical data were collected, and genetic analysis was performed on blood samples. Randomized selection ensured unbiased sampling, while laboratory personnel were blinded to case-control status to minimize bias. Genotyping analysis was conducted at the Rufaida BioMeds Laboratory in Dhaka, Bangladesh. To ensure participant confidentiality, the authors had no access to personally identifiable information during or after the data collection.

### 2.2. DNA Isolation & Genotyping

The blood samples were collected from all the participants and filled in the EDTA containing tubes. For the SNP analysis genomic DNA was extracted from peripheral blood leukocytes with the help of commercially available genomic DNA extraction kit (Favorgen, USA) following the given protocol. In brief, blood samples were lysed and then transferred into the binding column tubes with utmost care. The collected DNA was cleaned with the help of various buffers. Then DNA was isolated by using nuclease-free water and stored at −20°C after confirmation on a 1% agarose gel (Lonza, USA). DNA concentration and purity were assessed using a UV-Vis spectrophotometer (Shimadzu Corporation, Japan), and samples with an A260/280 ratio between 1.8 and 2.0 were included for downstream analysis.

Later, a thermal cycler (miniPCR, USA) was used to perform the polymerase chain reaction-restriction fragment length polymorphism (PCR-RFLP) assay using different primers (Macrogen, Korea) and it was utilized to detect single nucleotide polymorphisms (SNPs) at different genomic sites. A summary of the thermal condition and primer details to amplify the target details are mentioned in Table 1. EmeraldAmp GT PCR Master Mix (TAKARA BIO INC, Japan) helped to perform PCR and 1% agarose gel electrophoresis confirmed the completion of the PCR. Soon after completion of PCR, *MspI* and *BsrI* restriction enzymes (TAKARA BIO INC, Japan) (at 60°C for 1 hour in both cases) were used to digest the products. *MspI* and *BsrI* restriction enzymes used to examine the SNP at rs2279744 & rs937282, respectively. All the digested products were fractionated electrophoretically on 3% agarose gel with Midori Green (Nippon Genetics, Europe). All gels were independently reviewed by two analysts; discrepancies were resolved by repeat amplification and digestion. To validate the reliability of the data, 20% of the samples were subjected to Sanger sequencing through a commercial service provider, concordance with PCR-RFLP was 100%.

**Table 1.**
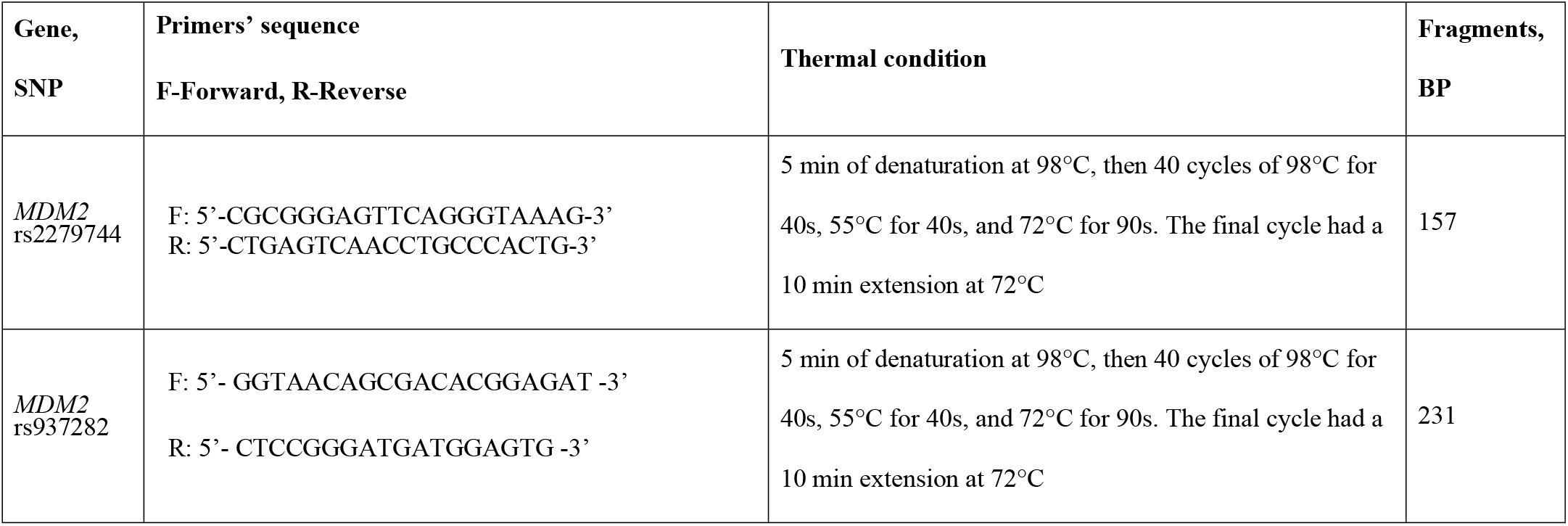
Primer sequences, thermal conditions and amplified products’ size.

### 2.3. Measurement of Serum *MDM2 & p53* Level

Circulating levels of *MDM2* and *p53* were quantified in serum using commercially available sandwich ELISA kits according to the manufacturers’ protocols. *MDM2* levels were measured using the Human *MDM2* ELISA Kit (catalog no. RAB1578, Sigma-Aldrich, Darmstadt, Germany), and *p53* levels were determined using the *TP53* ELISA Pair Set (catalog no. SEK90001, Sino Biological, Beijing, China), which detects full-length *p53* protein. For both assays, all reagents and samples were brought to room temperature before use, and all measurements were performed in duplicate.

For the *MDM2* assay, serum samples were diluted twofold using assay diluent, and 100 μL of diluted samples or serially diluted standards were added to the wells and incubated for 2.5 hours at room temperature with gentle shaking. After washing, biotinylated detection antibody was added and incubated for 1 hour. Then samples were kept with horseradish peroxidase–conjugated streptavidin for 45 minutes at room temperature. Plates were then washed and developed using TMB substrate for 30 minutes in the dark, and absorbance was measured immediately at 450 nm using 96 well plate spectrophotometer (BIOBASE, China).

For the *p53* assay, microplates were coated overnight at 4°C with capture antibody and subsequently blocked for 1 hour at room temperature. Serum samples were diluted fourfold with sample dilution buffer, and 100 μL of diluted samples or standards were added to the wells and incubated for 2 hours at room temperature. After washing, detection antibody was applied for 1 hour, followed by substrate incubation for 30 minutes in the dark. The reaction was stopped, and absorbance was read at 450 nm.

For both assays, standard curves were generated using a four-parameter logistic regression model (R^2^ ≥ 0.99). Concentrations of *MDM2* and *p53* in serum samples were calculated by interpolation from the corresponding standard curves and adjusted for sample dilution factors using GraphPad Prism 10.

### 2.4. Statistical Analysis

Statistical analyses were performed using R (version 4.4.1) and GraphPad Prism (version 9/10). Genotype distributions were tested for Hardy–Weinberg equilibrium using the HardyWeinberg package. Categorical variables were compared using Pearson’s chi-square test. Associations between *MDM2* polymorphisms (rs2279744 and rs937282) and BC risk were evaluated using multivariate logistic regression under additive, dominant, recessive, and over-dominant models, with age and BMI included as covariates. Adjusted odds ratios (aORs) with 95% confidence intervals (CIs) were estimated.

Because circulating biomarker data were non-normally distributed, non-parametric tests were applied. Serum *MDM2* and *p53* levels were compared between cases and controls using the Mann– Whitney U test. Genotype- and receptor subtype–dependent differences were analyzed using the Kruskal–Wallis test followed by Dunn’s multiple comparisons test. The association between circulating *MDM2* and *p53* levels was assessed using Spearman’s rank correlation among BC cases. Post hoc power for primary genetic models was estimated using observed control genotype frequencies with a two-sided α of 0.05. A p-value < 0.05 was considered statistically significant.

### 2.5. Ethical Consideration

The study received approval from the Institutional Review Board (IRB) of BRAC University (Approval No. BRACUIRB120220005). Written informed consent was obtained from all participants, who were assured of their right to withdraw from the study at any time without consequences. All procedures adhered to the principles of the Declaration of Helsinki. This study was reported in accordance with STREGA guidelines for genetic association studies.

## 3. Results

### 3.1. Socio-demographic and Clinical Characteristics

Socio-demographic and clinical characteristics are summarized in Supplementary Tables S1 and S2. Cases and controls were comparable across key variables, including age, BMI, and other socio-demographic factors (all p > 0.05), indicating well-matched groups.

Among patients, invasive ductal carcinoma was the predominant subtype. Most tumors were of moderate size, and hormone receptor positivity was common, with a subset showing lymph node involvement.

### 3.2. Serum *MDM2* and *p53* Levels in BC and HCs

Circulating levels of *MDM2* and *p53* were significantly altered in BC patients compared with HCs (Figure 1). Serum *MDM2* concentrations were markedly elevated in BC patients, whereas circulating *p53* levels were significantly reduced (both p < 0.0001, Mann–Whitney U test). These reciprocal changes indicate dysregulation of the *MDM2*–*p53* association in BC (Figure 1A–B).

**Fig. 1.**
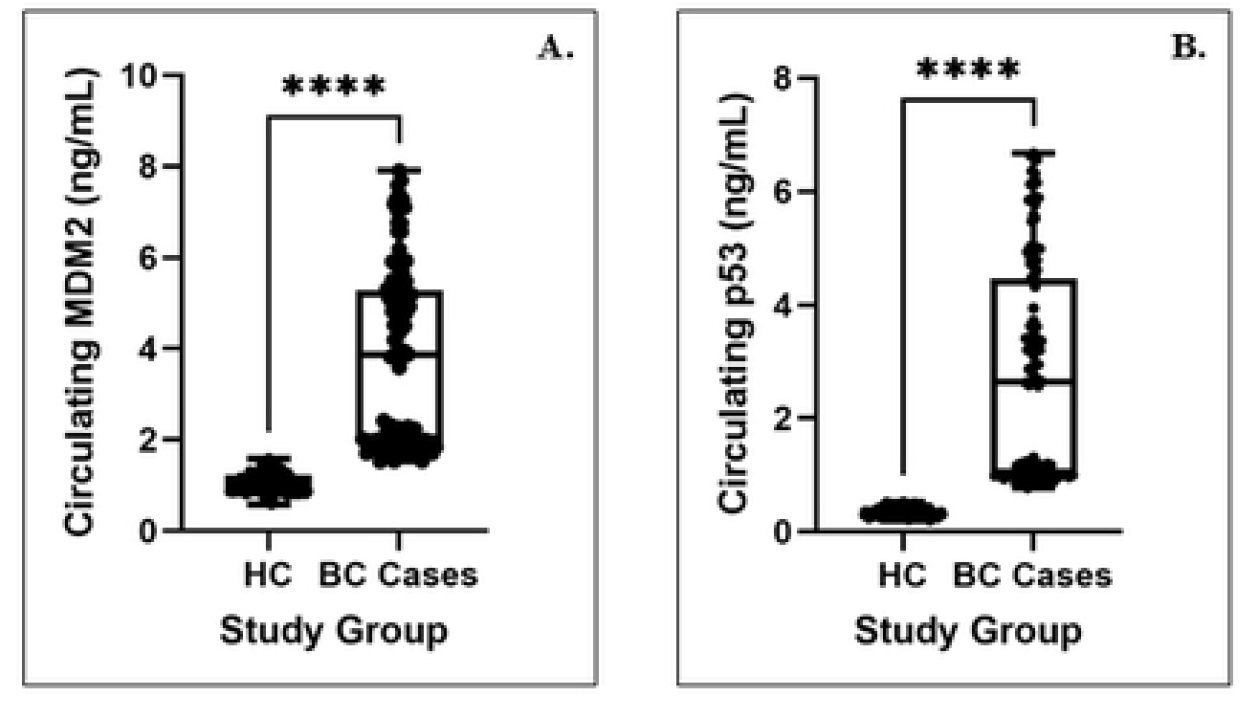

### 3.3. Association of *MDM2* rs2279744 Polymorphism with BC Risk

Genotype distributions of *MDM2* rs2279744 among cases and controls conformed to Hardy– Weinberg equilibrium (p > 0.05; Table 2). Multivariate logistic regression analysis adjusted for age and BMI revealed a strong association between rs2279744 and BC risk.

**Table 2.** Association between BC with *MDM2* rs2279744 Polymorphism (Multivariate Logistic Regression Adjusted for Age and BMI)

| Allele | Cases<br>(n = 112) | HWE |  | Controls<br>(n = 124) | HWE |  | Genetic models | aOR | 95% CI | P-value |
| --- | --- | --- | --- | --- | --- | --- | --- | --- | --- | --- |
|  |  | X<sup>2</sup> | P |  | X<sup>2</sup> | P |  |  |  |  |
| TT<br>Homozygote (157 bp) | 10<br>(8.93%) | 0.01 | 0.92 | 33<br>(26.613%) | 0.479 | 0.489 | Additive model 1<br>(TG vs. TT) | 11.258 | 4.621 – 27.427 | <0.001 |
|  |  |  |  |  |  |  | Additive model 2<br>(GG vs. TT) | 1.962 | 0.882 – 4.366 | 0.0986 |
| TG<br>Heterozygote<br>(157 bp, 109 bp and<br>48 bp) | 58<br>(51.79%) |  |  | 17<br>(13.710%) |  |  | Dominant model<br>(TG+GG vs. TT) | 3.699 | 1.727 – 7.924 | <0.001 |
| GG<br>Homozygote<br>(110 bp and 24 bp) | 44<br>(39.29%) |  |  | 74<br>(59.677%) |  |  | Recessive model<br>(GG vs. TT+TG) | 0.437 | 0.259 - 0.737 | 0.0018 |
|  |  |  |  |  |  |  | Over dominant model<br>(TG vs. TT+GG) | 6.760 | 3.594 – 12.717 | <0.001 |

**Note:** Genotype frequencies were tested for Hardy-Weinberg equilibrium (HWE) using the Chi-square test (X<sup>2</sup>). Adjusted Odds Ratios (aOR), 95% Confidence Intervals (CI), and p-values were calculated using multivariate logistic regression in R. All models used the TT genotype (common homozygote) as the reference group.
p > 0.05 indicates that genotype distribution follows Hardy-Weinberg equilibrium.
Abbreviations: HWE – Hardy-Weinberg equilibrium, aOR – Adjusted Odds Ratio, CI – Confidence Interval, X<sup>2</sup> – Chi-square value

A significant association between rs2279744 and BC risk was observed. In the additive model, women carrying the heterozygous TG genotype had a markedly increased risk of BC compared with TT carriers (aOR = 11.258, 95% CI: 4.621–27.427, p < 0.001). In contrast, the GG genotype did not show a significant increase in risk in the additive model (aOR = 1.962, 95% CI: 0.882– 4.366, p = 0.099).

In the dominant model, carriers of at least one G allele (TG+GG) had a significantly higher risk of BC than TT carriers (aOR = 3.699, 95% CI: 1.727–7.924, p < 0.001). Notably, the over-dominant model showed that the TG genotype alone conferred a substantially increased risk when compared with both homozygous genotypes combined (aOR = 6.760, 95% CI: 3.594– 12.717, p < 0.001).

In contrast, the recessive model indicated a protective effect of the GG genotype (aOR = 0.437, 95% CI: 0.259–0.737, p = 0.001). Together, these results suggest a genotype-dependent and non-linear effect of rs2279744 on BC susceptibility (Figure 2A; Table 2).

**Fig. 2.**
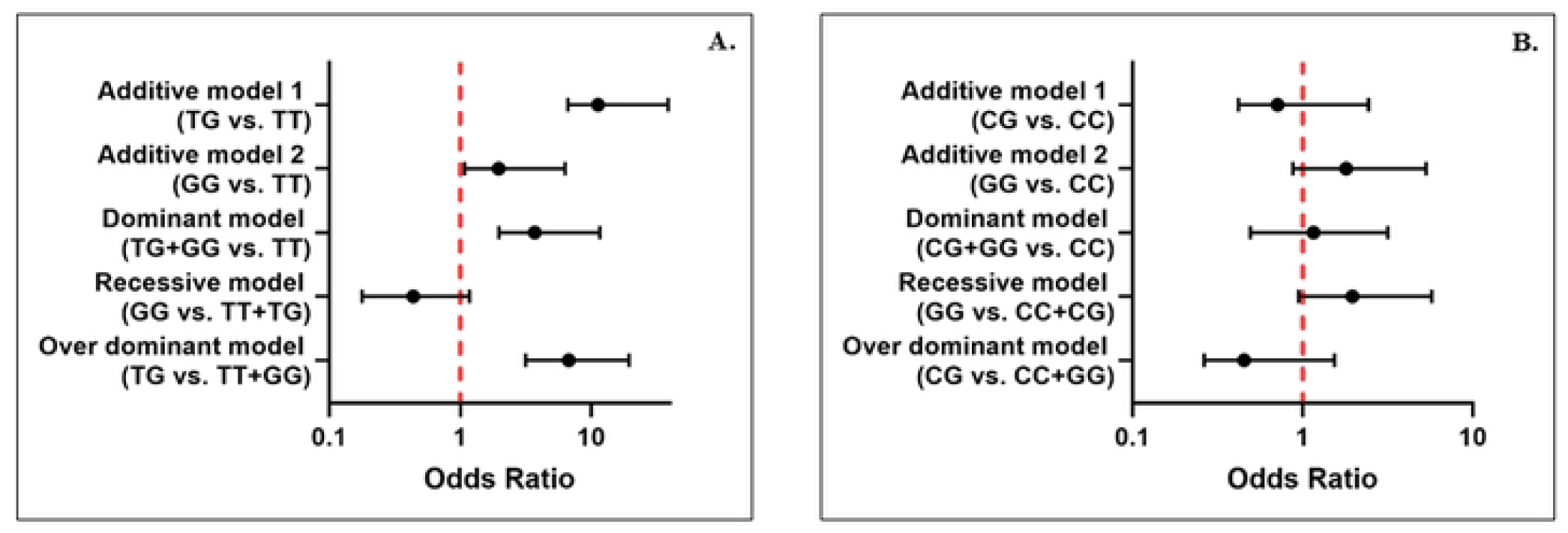

### 3.4. Association of *MDM2* rs937282 Polymorphism with BC Risk

Genotype frequencies of *MDM2* rs937282 also followed Hardy–Weinberg equilibrium in both groups (Table 3). Unlike rs2279744, rs937282 showed no strong association with BC risk across most genetic models.

**Table 3.**
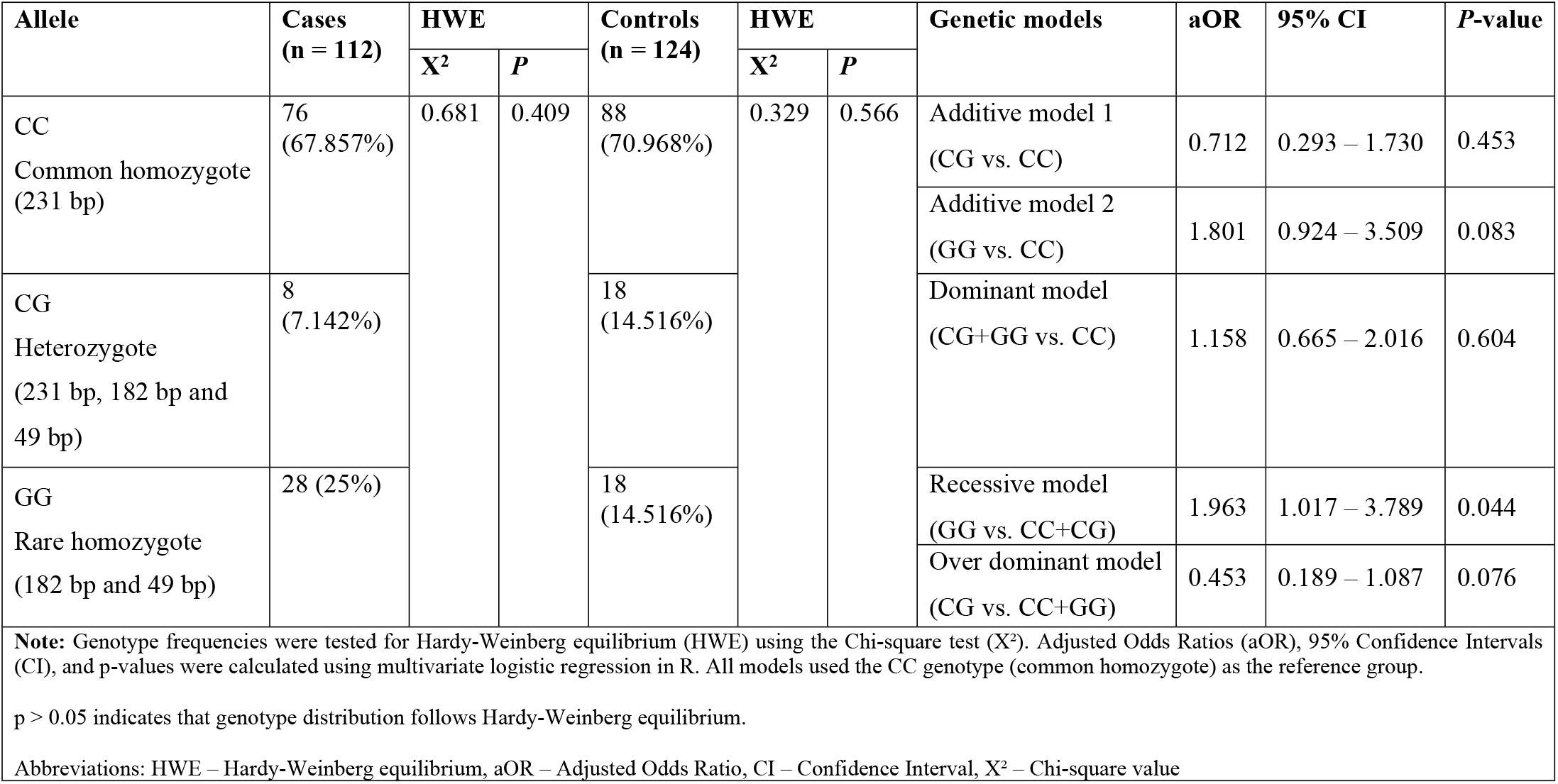
Association between BC with *MDM2* rs937282 polymorphism (Multivariate Logistic Regression Adjusted for Age and BMI)

In the additive model, the CG genotype did not show a significant association with BC risk compared to the CC genotype (aOR = 0.712, 95% CI = 0.293–1.730, p = 0.453). The GG genotype, however, showed a trend toward increased risk (aOR = 1.801, 95% CI = 0.924–3.509, p = 0.083), though it did not reach statistical significance.

In the dominant model (CG + GG vs. CC), no significant association was observed (aOR = 1.158, 95% CI = 0.665–2.016, p = 0.604). Under the recessive model, the GG genotype showed a borderline association with increased risk (aOR = 1.963, 95% CI: 1.017–3.789, p = 0.044); however, given the limited statistical power and small number of GG carriers, this finding should be interpreted with caution. The over-dominant model suggested a possible protective effect of the CG genotype (aOR = 0.453, 95% CI: 0.189–1.087, p = 0.076), although this association was not statistically significant. Overall, none of these associations remained robust, indicating a limited role of rs937282 in BC susceptibility in this cohort (Figure 2B; Table 3).

### 3.5. Genotype-Dependent Variation in Circulating *MDM2* and *p53* Levels

Serum *MDM2* and *p53* concentrations differed significantly across rs2279744 genotypes (Figure 3). TG carriers exhibited the highest circulating *MDM2* levels, whereas TT carriers showed the lowest levels. Conversely, circulating *p53* concentrations were significantly reduced in TG carriers compared with TT and GG genotypes (p < 0.0001). These findings suggest that the rs2279744 polymorphism exerts functional effects on the *MDM2*–*p53* regulatory association (Figure 3A–B).

**Fig. 3.**
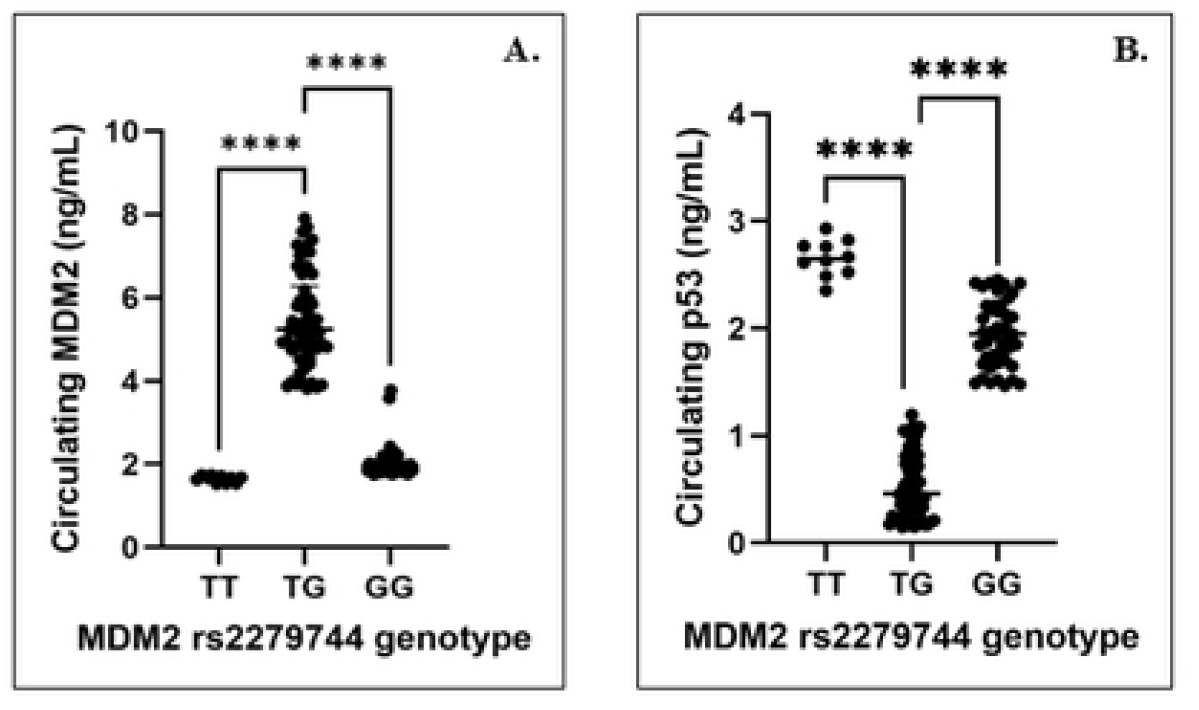

### 3.6. Correlation Between Circulating *MDM2* and *p53*

Spearman’s rank correlation analysis revealed a strong inverse relationship between circulating *MDM2* and *p53* levels (r = −0.78, p < 0.0001). Higher *MDM2* concentrations were associated with lower *p53* levels, supporting enhanced *p53* degradation in BC patients (Figure 4).

**Fig. 4.**
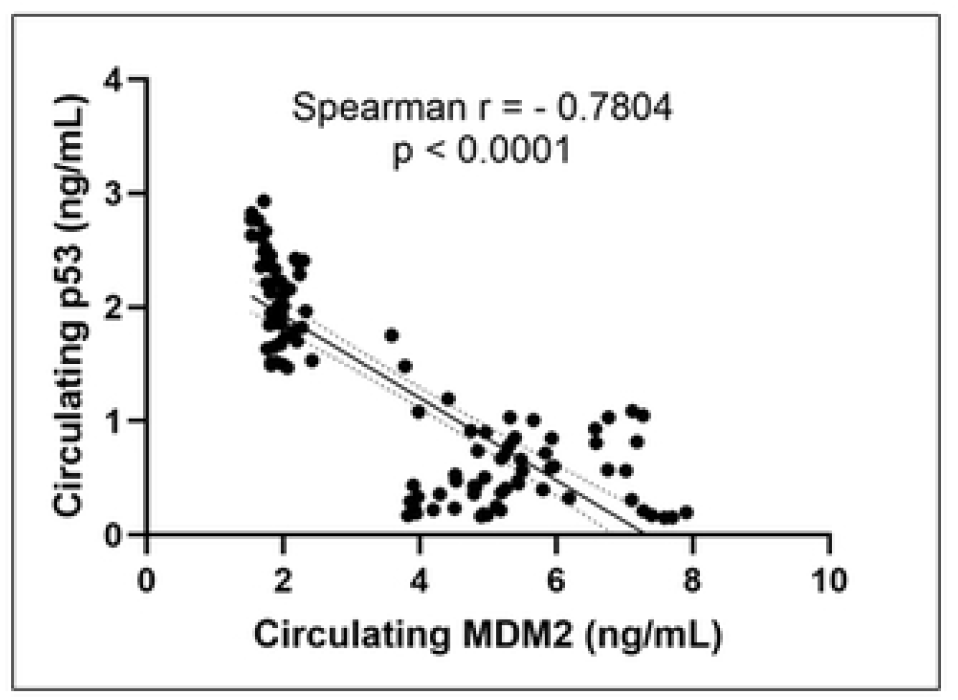

### 3.7. Circulating *MDM2* Levels Across BC Receptor Subtypes

Circulating *MDM2* levels varied significantly across BC receptor subtypes (Figure 5). Overall group differences were significant (Kruskal–Wallis p < 0.0001). *HER2*-positive tumors and triple-positive (*ER/PR/HER2*-positive) tumors exhibited the highest circulating *MDM2* levels, whereas *HER2*-negative tumors showed the lowest levels. Compact letter display analysis confirmed statistically distinct subgroup patterns following Dunn’s multiple comparisons test (Figure 5). Overall, the combined genetic and protein expression analyses support a genotype-dependent disruption of the *MDM2* regulatory pathway in BC.

**Fig. 5.**
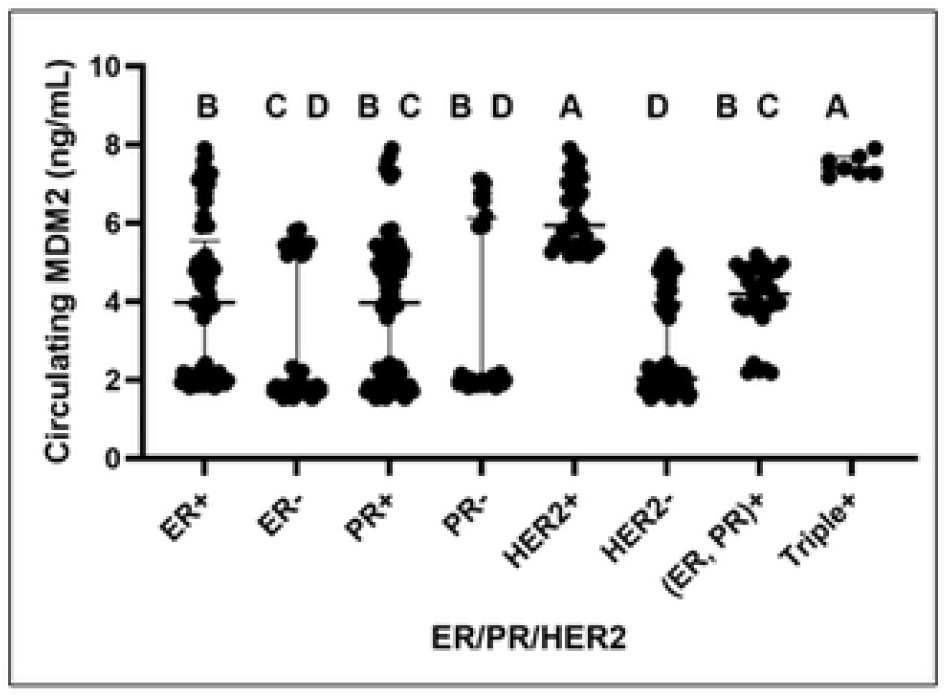

## 4. Discussion

This study analyzed the association of two regulatory polymorphisms in the *MDM2* gene, rs2279744 and rs937282, with BC risk in 112 cases and 124 HCs. Based on an assumed minor allele frequency of approximately 0.30 and a significance level of α = 0.05, the estimated statistical power exceeded 95% for rs2279744 due to its moderate and consistent effect sizes. In contrast, rs937282 demonstrated smaller effect sizes, with estimated power ranging from 55–70%. Thus, while the sample size was sufficient to detect strong genetic associations, weaker effects; particularly for rs937282, may require confirmation in larger cohorts.

The primary finding of this study is a significant and genotype-dependent correlation between rs2279744 and breast cancer susceptibility in Bangladeshi women, whereas rs937282 exhibited no statistically significant effect. The heightened risk was predominantly attributed to the heterozygous TG genotype of rs2279744, signifying a non-linear regulatory influence rather than a straightforward allele-dose effect. The lack of a clear link for rs937282 makes the rs2279744 finding more specific and less likely that the effect seen is due to population stratification or methodological bias.

*MDM2* is an important negative regulator of the tumor suppressor protein *p53*, which stops cells from growing out of control by controlling the cell cycle, fixing damaged DNA, and causing cell death when necessary [35]. Genetic variants that modify *MDM2* transcription can disrupt the *p53– MDM2* feedback loop, resulting in diminished *p53* activity and enhanced survival of cells with genomic damage [36, 37]. This biological framework provides strong support for investigating regulatory *MDM2* variants in cancer susceptibility.

The rs2279744 polymorphism (SNP309, T>G) is located in the promoter region of *MDM2* and has been shown to enhance binding of the transcription factor Sp1, resulting in increased *MDM2* expression [36]. In the present study, carriers of the TG genotype showed a significantly increased risk of BC across additive, dominant, and over-dominant genetic models. In contrast, the GG genotype did not show a significant increase in risk in additive models and appeared to be protective under the recessive model. This pattern indicates that the effect of rs2279744 is not driven simply by the presence of the G allele, but rather by genotype-specific regulatory dynamics.

Importantly, our circulating biomarker data supports this mechanistic interpretation. Women with TG had the highest levels of *MDM2* in their blood, but their levels of *p53* were lower. The strong negative correlation between circulating *MDM2* and *p53* makes it even more clear that this variant plays a functional role in changing the *MDM2–p53* association. Although circulating *p53* levels may reflect protein release or stabilization rather than fully active tumor suppressor function, their consistent inverse relationship with *MDM2* clearly indicates dysregulation of this pathway.

The finding that the TG genotype presents a greater risk compared to the GG genotype substantiates the concept of heterozygote-specific regulatory imbalance, as opposed to a linear allele-dose response. Even small increases in MDM2 expression could have a major impact on p53 function in the tightly controlled feedback loop [18]. Such non-linear regulatory effects may explain why heterozygous carriers exhibited the highest disease risk and *MDM2* levels, whereas homozygous GG carriers did not consistently show the same magnitude of effect. The small variation seen between the TT and TG genotypes suggests that there may be an allele-dose effect, but confirmation in larger groups would be valuable.

These results align with previous studies in other populations, where rs2279744 has been identified as a functional SNP that alters *MDM2* level [38]. The G allele has been linked to increased transcriptional activity, reduced *p53*-mediated apoptosis, and enhanced tumor development [39].

Studies in Chinese populations similarly reported significant associations for the TG genotype, although the effect of the GG genotype varied across genetic models [24]. Taken together, this suggests that carrying one copy of the G allele (heterozygosity) may have a particularly strong impact on disease risk, likely due to its influence on promoter activity and gene expression balance [40].

In contrast, the rs937282 polymorphism did not demonstrate a consistent association with BC risk in this study. Although the GG genotype suggested a nearly two-fold increased risk under the recessive model and the CG genotype appeared potentially protective under the over-dominant model, these findings were not robust across models. This inconsistency mirrors prior reports, where rs937282 has shown variable or modest associations depending on population and study design [41–43]. Located in the 5′UTR of *MDM2*, rs937282 may influence mRNA stability or translational efficiency [19, 41], but its regulatory effect appears less direct than the promoter-based impact of rs2279744 [44].

In addition to genetic associations, our examination of tumor subtypes indicated increased circulating *MDM2* levels in *HER2*-positive and triple-positive tumors. These results indicate possible interactions between oncogenic receptor signaling and *MDM2*-mediated inhibition of *p53. HER2*-driven tumors often act aggressively, so higher *MDM2* activity may help these types of tumors grow even more. So, *MDM2* dysregulation may be especially important in BC phenotypes that are biologically aggressive or driven by receptors.

From a clinical perspective, the tumor characteristics observed in our cohort align with global trends. The main subtype was invasive ductal carcinoma, and most of the tumors had hormone receptors, which means they might respond to endocrine therapies [45–48]. Approximately 29% of patients were *HER2*-positive, comparable to rates reported in other Asian populations [49, 50]. A lot of the tumors were medium to large, and a lot of them had lymph node involvement, which shows that Bangladesh needs better ways to find them early [51, 52].

Importantly, cases and controls were well matched across socio-demographic variables, including age, BMI, education, socio-economic status, reproductive history, and menopausal status. This demographic balance enhances the validity of the observed genetic associations and diminishes the potential for confounding effects.

In general, our results show that the two *MDM2* variants have very different effects on how likely someone is to get BC. The rs2279744 polymorphism looks like a biologically important regulatory variant that has clear functional and clinical significance. On the other hand, rs937282 has weaker and less consistent effects. These findings underscore the significance of examining genotype-specific regulatory dynamics within the *p53–MDM2* pathway and emphasize rs2279744 as a potential biomarker for breast cancer risk stratification in South Asian populations.

### Limitations and Future Directions

Several limitations should be acknowledged. First, the small sample size may have made it more difficult to find weaker genetic effects, especially for rs937282. Second, functional assays were not conducted to directly evaluate allele-specific *MDM2* expression or *p53* activity. Third, the design of the study may not be applicable to other settings because it was done in a single hospital. Future multicenter studies that include functional validation and long-term follow-up are necessary.

## 5. Conclusion

In summary, this research identifies *MDM2* rs2279744 as a significant genetic factor influencing breast cancer risk in Bangladeshi women, particularly highlighting the pronounced impact of the heterozygous TG genotype. This risk seems to be caused by problems with the *MDM2–p53* association, which is when *MDM2* levels are too high and *p53* levels are too low. Conversely, rs937282 exhibited no significant correlation with BC susceptibility. These results highlight the biological and clinical significance of regulatory *MDM2* variants and support for additional investigation of the *MDM2–p53* pathway as a prospective biomarker and therapeutic target in breast cancer.

## Ethics Statement

The study was approved by the Institutional Review Board (IRB) of BRAC University (Approval No. BRACUIRB120220005)

## Acknowledgement

We thank all the participants and their relatives for their cooperation in this study. Also, we would like to thank all physicians and administrative staff at the National Cancer Research Institute and Hospital, Mohakhali, Dhaka, Bangladesh, for their support in data and sample collection. Also, we would like to thank Rufaida BioMeds for kindly allowing us to use their research facilities.

## Author Contributions

Conceptualization: M.A.H.; Data curation: M.N.I., N.B.H., M.I.S., and M.H.T.; Formal analysis: M.N.I., A.A.K., J.T., and S.M.; Investigation: M.H.C., F.I., A.A.K., and M.A.S.; Methodology: M.A.H., and M.H.C.; Supervision: M.A.H.; Writing – original draft: F.I.; Writing – review & editing: M.A.H.

## Data availability statement

Data available on request due to privacy/ethical restrictions.

